# Non-equilibrium processes prevail in shaping species richness and functional diversity of terrestrial vertebrates in a global hotspot

**DOI:** 10.1101/2022.12.19.521041

**Authors:** Matheus de Toledo Moroti, Alexander Skeels, Fernando R. Da Silva, Diogo B. Provete

## Abstract

**Aim:** The effects of equilibrium and nonequilibrium processes are generally investigated using species richness on a single biological group. However, little is known about how these two classes of processes also affect trait diversity, considering multiple taxa within the same geographical template. Here, we evaluated which variables representing equilibrium (topography, climate, and primary productivity) and nonequilibrium (diversification rate and evolutionary time) processes best explain species richness and trait diversity of four clades of vertebrates within the same global hotspot. We also investigated how trait disparity has accumulated over time and whether there are congruent spatial patterns between groups.

**Location:** Atlantic Rainforest

**Time period:** Contemporary.

**Major taxa studied:** Terrestrial vertebrates.

**Methods:** We tested whether the spatial pattern of Functional Dispersion (FDis), richness, diversification rate, and evolutionary time of each group are correlated. We used a spatially explicit structural equation model to test how species richness and functional dispersion are influenced by variables representing equilibrium and nonequilibrium mechanisms. Additionally, we explored how trait disparity accumulated over time in the four groups.

**Results:** We found that non-equilibrium proxies, evolutionary time and diversification rate, played a primary role in driving species richness and trait diversity, with elevation and climate variables having only an indirect effect species and trait diversity via diversification rate and evolutionary time. We found a congruent pattern of species richness among all groups, except among ectotherms. In contrast, the spatial distribution of evolutionary time was distinct for each group.

**Main conclusions:** Despite nonequilibrium processes were more important for generating large-scale diversity patterns within the same geographical template, the interplay between evolutionary time and dispersal ability have disparately determined the assembly of communities.

## Introduction

Whether broad-scale geographical patterns of biological diversity are governed by environmentally determined carrying capacities (equilibrium dynamics; Rabosky & Hurlbert, 2015) or instead reflect historical differences in occupation times or diversification rates (non-equilibrium dynamics; Harmon & Harrison, 2015) is a fundamental question in biodiversity studies. Under an equilibrium model, features of the environment, and specifically measures of resource availability (such as net primary productivity) or resource diversity (such as environmental heterogeneity), are expected to be the strongest predictors of species diversity. This is because more resources, or kinds of resources, can facilitate differentiation of resource-use between species, allowing more species to coexist (Macarthur and Levins 1967; MacArthur 1972). Yet regions do not only differ in present-day measures of resource diversity and availability, they also differ in their geological and environmental histories, which may determine the presence or absence of factors driving speciation, extinction, and colonization dynamics through deep-time. Therefore, under a non-equilibrium model, species diversity within regions can also be the outcome of the historical dynamics that shape the accumulation of species relative to other regions (Rohde 1992). Studies investigating the relative support for these two models in explaining present-day biodiversity gradients have shown mixed results. However, most studies investigating the relative importance of these processes are based on single clades and focus exclusively on species diversity. Consequently, it is unclear whether equilibrium and non-equilibrium processes have the same importance explaining the variation in diversity for ecologically and evolutionarily distinct clades that share similar geological and environmental selective pressures within the same geographical region.

The relative importance of equilibrium vs. non-equilibrium processes may vary depending on biological characteristics that determine resource use and interactions with the abiotic environment, including metabolic rates, thermoregulatory modes, and dispersal ability (Buckley *et al.*, 2012). In ectotherms, activity time is limited by available environmental energy, while endotherms can remain highly active if resource consumption meets their metabolic demands (Buckley *et al.*, 2012). As a result, endotherms require more resources than ectotherms to thermoregulate, particularly in colder environments (Clarke & Gaston, 2006). Ectotherms also typically have narrower environmental tolerances and more selective habitat preferences, which leads to slower rates of environmental niche evolution, smaller geographic ranges, and narrower latitudinal and elevational ranges (Rolland *et al.*, 2018). This could make ectotherms more susceptible to environmental stochasticity and more dependent on climatic refuges (Carnaval *et al.*, 2009), which would increase rates of extinction though range collapse, or increase rates of speciation through range fragmentation, leading to different patterns of species distributions (e.g., Schweizer *et al.*, 2014; MacDougall *et al.*, 2019). Finally, ectotherms generally have lower dispersal capacity, which in combination with narrow environmental preferences, could mean that these species are less able to colonise new regions or novel habitats through time. Therefore, fundamental metabolic and dispersal constraints between ectotherms and endotherms may drive differences in resource-use and niche partitioning under an equilibrium model and rates of diversification and/or occupation times under a non-equilibrium model

In a geographic area, where historical and contemporary variations related to geology and climate conditions act as a geographic template for all biological groups, it is possible to detect how species and trait diversities of ectotherms and endotherms respond to equilibrium and non-equilibrium processes. The Brazilian Atlantic Forest harbours an outstanding diversity and endemism of terrestrial vertebrate species (Figueiredo *et al.*, 2021). Due to its wide latitudinal (from 5 to 33° S) and longitudinal (from 35 to 55° W) ranges, the Atlantic Forest is a heterogeneous environment that varies in climatic gradients, such as annual rainfall (from 800 to 4,000 mm), mean annual temperature (from 15 to 25 °C), and topography (from sea level to 2,200 m a.s.l.). Species richness in amphibians, terrestrial mammals, and birds peak in the highly productive central and coastal portion of the Atlantic Forest, especially in the Serra do Mar region (Vasconcelos et al. 2019). This suggests that environmental productivity could be a major determinant of species diversity. However, while reptiles also present greater richness in this region, they are closer to the region of contact with dry savannas in the inner part of the continent (Figueiredo *et al.*, 2021). In addition to present-day climatic differences, historical climate change was important in shaping species distributions in the regions. For example, phylogeographic studies have found high genetic diversity in frogs (Carnaval *et al.*, 2009), birds and lizards in the central region of the Atlantic Forest (reviewed in Peres et al. 2020), which was more climatically stable during the Pleistocene. Further, there is evidence of recent colonization and population expansion in areas that were less stable in the past, such as the southern portion (reviewed in Peres et al. 2020), indicating that past climate conditions might be associated with species geographic range expansion and extinction, thereby regulating the size and composition of regional species pools (Benício *et al.*, 2021). Thus, because of its exceptional biodiversity, sharp climatic gradients, and dynamic climatic history, the Atlantic Forest is an excellent model to understand the role of equilibrium vs. non-equilibrium processes in generating and maintaining spatial distribution patterns of multiple components of diversity for different taxa within the same geographical template.

Here, we evaluated the relative importance and spatial congruence of variables representing equilibrium and non-equilibrium processes explaining variation in species richness and trait diversity of vertebrates in the Atlantic Forest. Specifically, we tested (1) how species richness and trait diversity are related to evolutionary time, diversification rate, productivity, climate, and topographical heterogeneity (Figure 1). Under an equilibrium model we predict that species or trait diversity will be positively, and directly, related to climate, productivity or topographical heterogeneity, as these factors set the ecological limits on biodiversity (Rabosky & Hurlbert, 2015). As an equilibrium model predicts that high diversity is achieved via niche partitioning in high resource environments, we additionally predict that trait diversity (as a proxy for ecological diversity) should be positively related to species richness. Under a non-equilibrium model, we predict that species and trait diversity will be directly and positively related to diversification rates or evolutionary time (Harmon & Harrison 2015), and that topography, productivity or climate will indirectly drive species and trait diversity via their effect on diversification rates or evolutionary time. This could be because topography and climate can shape rates of speciation through allopatric divergence, or increase rates of extinction through climate change. Furthermore, as different biomes have different ages (Jetz and Fine 2012), and different regions have acted as climatic refugia through time, these variables might also reflect the age and stability of the assemblage. In addition, we tested (2) whether the distribution of vertebrate communities is congruent in terms of diversification rate, assemblage age, trait diversity, and species richness. Considering that amphibians and reptiles have narrower physiological requirements and lower dispersal ability than endotherms, which are usually better trackers of ecological limits than ectotherms (e.g., Araújo & Pearson, 2005; Rolland et al., 2018), we expect more similarity between ectotherms than between endotherms.

**Figure 1.**
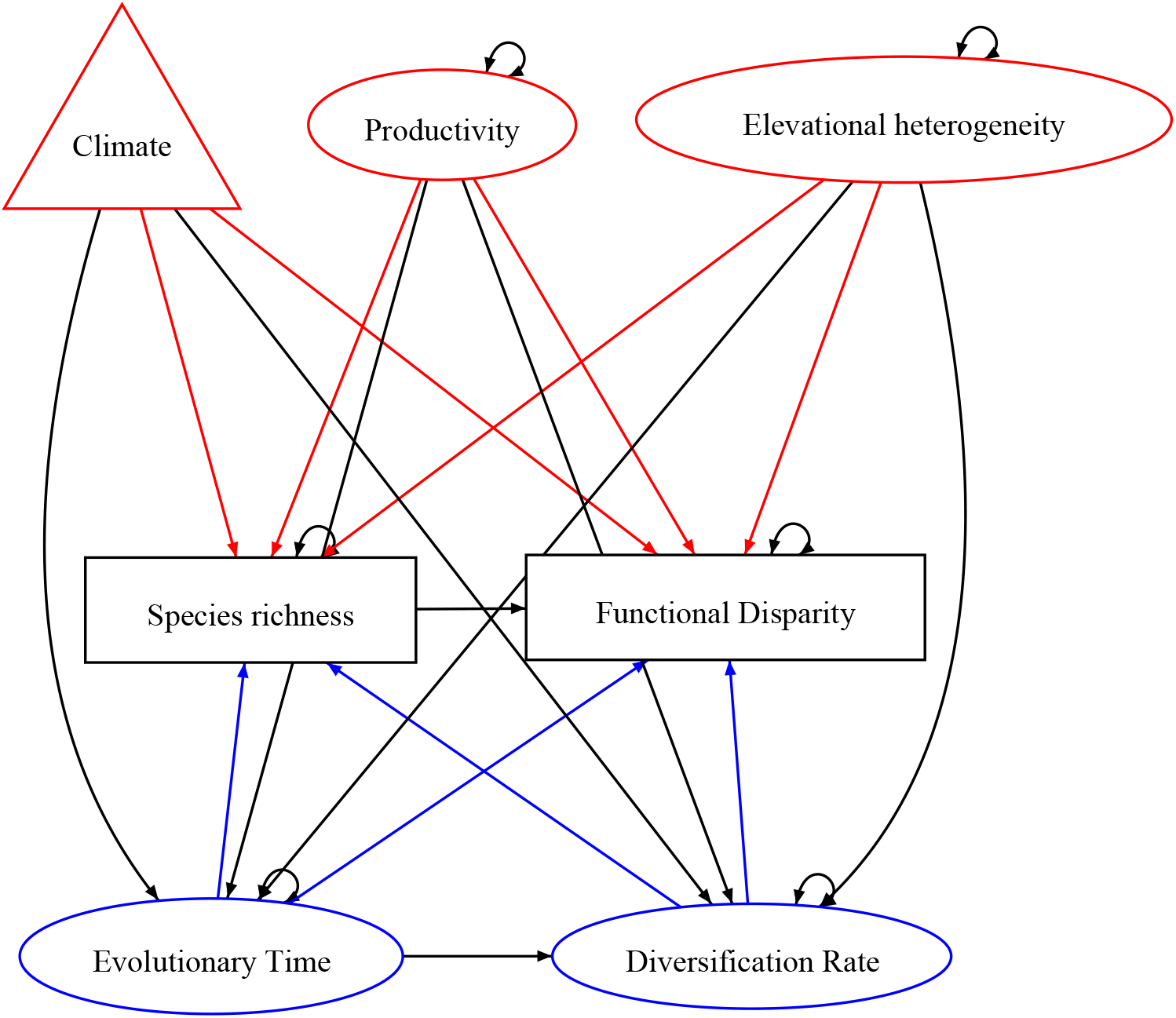
Hypothesis tested in this paper referring to each class of mechanism. The pathway connecting productivity to species richness represents the more individuals hypothesis, while the pathways connecting elevational heterogeneity/climate/productivity to functional dispersion reflect ecological niche partitioning. Red colour indicate non-equilibrium mechanisms, while blue represent equilirbium mechanisms. Triangle represents the statistical composite variable.

## Material and Methods

### Species geographical data

We built a 10 x 10 km grid with 0.5° cells overlaid on the Atlantic Forest consensus extent (Muylaert *et al.* 2018) in QGIS software. We excluded cells with less than 50% continental coverage, resulting in 424 cells. We obtained extent-of-occurrence polygons of vertebrates. Although these polygons are scale dependent (Hulbert and White, 2005) and include commission errors, recent studies (e.g., Silva *et al.*, 2016) have found congruence between point occurrence data and range maps for 10 x 10 Km scales for amphibians in the Atlantic Forest, whose distribution is less known than other taxa. For amphibians, we used data from IUCN (2021), complemented by shapefiles from (Vasconcelos *et al.*, 2019). For squamates, we used data from Roll *et al.* (2017). For birds, we used data from BirdLife (2015). For mammals, we used data from IUCN (2021). We obtained presence-absence matrices separately for each group by overlapping polygons on the grid using the R package *letsR,* considering that at least 50% of the polygon covered the cell. We defined our species pool as native and non-migrating species for the whole Atlantic Forest. Finally, our presence-absence matrices contained 518 species of frogs, 402 squamates, 815 birds, and 236 mammals. Species nomenclature followed the phylogeny of each group.

### Trait data

We used four response traits to describe the trait space of assemblages at large spatial scales: Body mass for endotherms and body size for ectotherms, Diel activity, and Habitat (Table S1). We chose these traits because they reflect different niche dimensions, in addition to being directly related to climate gradients (e.g. Oliveira *et al.*, 2016; Vasconcelos *et al.*, 2019). For example, body size is directly related to metabolic capacity in ectotherms and endotherms. Likewise, diel activity affects foraging patterns in birds and mammals, and is related to risk desiccation in ectotherms.

We obtained data on habitat, body size, and diel activity from the following sources: i) amphiBIO (Oliveira *et al.*, 2017) and Vasconcelos *et al.* (2019) for frogs; ii) Marques et al. (2019) for Squamates; iii) Elton Traits (Wilman *et al.*, 2014) for birds and mammals, and complement with PanTHERIA for mammals. We also complemented them with the literature for 9% of frogs, 22% of squamates, and 10% of mammals (Supplementary material). However, 84 frog (~17%), 135 squamate (~34%), 107 bird (~13%), and 41 mammal species (~17%) had missing trait values. Thus, they were excluded from the analyses, resulting in a database with 434, 267, 708, and 195 species, respectively.

Missing trait data is expected in global databases and can lead to bias in functional metrics. Although there are imputation methods for continuous traits with good performance, there is still a need to explore methods of imputation and errors with categorical traits. In addition, imputation methods still have a bias with high missing data values (>40%), which can only be better resolved with field data collection and natural history museums (Johnson *et al.*, 2020).

### Phylogeny

We used the consensus tree for amphibian containing 7,239 species of Jetz & Pyron (2018). For squamates, we used the fully-sampled consensus tree with 9,754 species of Tonini *et al.* (2016). For birds, we used the tree proposed by Jetz *et al.* (2012), with 9,993 bird species. For mammals, we used the fully-sampled tree of Upham *et al.* (2019). Birds are the only group to which a consensus tree is not provided. Thus, following Rubolini *et al.* (2015), we generated 10,000 posterior trees to our species pool based on the ‘Hackett’ backbone (Jetz *et al.*, 2012). Then, we built a Majority Rule Consensus tree with the *DendroPy* and *SumTrees* packages in Python.

We pruned the species to which we had occurrence and trait data from the respective phylogenies using the R package *picante.* Yet, 24 frog (~3%), 70 birds (~10%), and 28 mammals (~14%) species had not been sampled in the phylogenies and were removed from analysis. Thus, our final dataset contained 421 frog, 261 squamates, 638 birds, and 167 non-flying mammal species. Our final dataset corresponds to ~60% of amphibian richness (707 species), ~53% of reptiles (492 species), 62.2% of birds (1025 species), and ~64% of mammals (262 species) known to occur in the Atlantic Forest (Figueiredo *et al.*, 2021).

### Trait Diversity

To describe the multidimensional trait space, we calculated Functional Dispersion (FDis; Laliberté & Legendre, 2010) using the trait distance and presence-absence matrices in the FD package. We chose this metric because it can identify areas with the largest differences in trait diversity (Fig. S1, S2). FDis is the average Euclidean distance of species to the centroid of the functional volume (Laliberté & Legendre, 2010). Therefore, the metric measures the size of the functional space of the community (Mammola *et al.*, 2021), independently from species richness. Simulation studies (Kuebbing et al. 2017, McPherson et al. 2018) have shown that FDis has a good performance to detect community assembly mechanisms. It is also not prone to be influenced by outliers. To calculate the trait dissimilarity matrix, we used the Gower index that allows for mixed data (Pavoine *et al.*, 2009) in the R package *ade4.*

### Variables related to equilibrium processes

Equilibrium processes assume that environmental conditions (e.g., productivity, climate, area), as surrogates for resource availability, determine the ecological limits and opportunities for niche partitioning that dictate the maximum number of species a region can support (Rabosky & Hurlbert, 2015). Therefore, species richness and composition would result from the evolution of ecological differences that determine the ability of species to persist in the environment over time (Rabosky, 2013). We used three environmental variables that reflect the total pool of resources available to represent equilibrium processes in our structural equation model (SEM): i) Productivity – high productivity and resource availability can sustain more individuals, more viable populations, and more species (i.e., the “more individuals” hypothesis; Brown, 1981). For each grid cell, we calculated the mean and the coefficient of variation of net primary productivity obtained from the MOD17A2H variable (Running, 2013); ii) Topographical heterogeneity – altitudinal variation affords a wide diversity of niches, facilitating species coexistence. We obtained the coefficient of variation of elevation at 1 km from Amatulli *et al.*, (2018), and iii) climate – climatic variables determine range limits in vertebrates (e.g., Wiens *et al.*, 2006; Botero *et al.*, 2014), and potentially influence large-scale species distributions. We firstly chose seven bioclimatic variables from (Karger & Zimmermann, 2019): Mean Diurnal Temperature Range (BIO2), Isothermality (BIO3), Temperature Seasonality (BIO4), Temperature Annual Range (BIO7), Mean Temperature of Coldest Quarter (BIO11), Precipitation Seasonality (BIO15), and Precipitation of Driest Quarter (BIO17). After testing for multicollinearity, we excluded BIO2 and BIO17 that had Variation Inflation Factor >10 from further analysis.

To reduce model complexity, BIO3, BIO4, BIO7, BIO11, and BIO15 were further combined into a statistical composite variable (Grace & Bollen, 2008; Grace et al. 2010) called ‘climate’. The variable with the highest loading is BIO3. Positive loadings mean a more stable climate. This analysis was performed in the R software (R Core Team, 2020).

### Variables related to non-equilibrium processes

Non-equilibrium processes assume that species richness on a continental scale would be determined solely by colonization time (i.e., time-for-speciation effect, Wiens, 2011), diversification rate, and subject to stochastic dynamics that constantly change the limit of species (Harmon & Harrison, 2015). We calculated two predictor variables to represent non-equilibrium processes separately for each group: i) Mean Pairwise Distance (MPD) to describe assemblage age (Fig. S7) (Webb *et al.*, 2002) – MPD increases with the degree of phylogenetic divergence between species within a community, which makes it a proxy for evolutionary time (e.g., Oliveira *et al.*, 2016). Oliveira *et al.* (2016) using simulations found that MPD was the best metric to describe assemblage age in mammals worldwide. Therefore, MPD could be used to indicate non-equilibrium mechanisms if there is an accumulation of species in areas with higher MPD (time-for-speciation effect, Wiens, 2011), or if areas with higher functional dispersion also have higher MPD, which would indicate a longer evolutionary time to overcome niche conservatism (e.g., Oliveira *et al.*, 2016); ii) diversification rates – High diversity might result from a fast accumulation of species due to high speciation and/or low extinction. We calculated the tip-level diversification rate (DR; Jetz *et al.*, 2012) in the *picante* package. This method is the inverse of the Equal Splits measure (Redding & Mooers, 2006) and allows calculating the diversification rate for each branch of the phylogeny. Rapidly diversifying lineages have high DR values (Jetz *et al.*, 2012). Subsequently, we calculated the average of the DR for each grid cell (Figure S4).

There is no consensus on how accurate the speciation rate estimated with these metrics is, thus biasing diversification measures towards speciation and down-weighting extinction (Title & Rabosky 2019). However, DR appears to be a suitable metric to answer our questions, as it is a species level metric, incorporates the number of split events, and distances between nodes from the root to tips of a phylogeny, while giving greater weight to branches closer to the present (Redding & Mooers, 2006; Jetz *et al.*, 2012).

### Data Analysis

To test whether there is spatial congruence in species richness, functional dispersion, MPD, and DR between the four groups, we used Tjøstheim (1978) pairwise correlation coefficient in the *SpatialPack* R package. This coefficient is a non-parametric analysis that tests the association between two spatial variables. Values above 0.46 indicate a positive spatial correlation.

We used spatially-explicit Structural Equation Models. For that, we built four Generalized Least Squares (GLS) models to test how each predictor variable (climatic, productivity, topographical heterogeneity, diversification rate, and evolutionary time) was related to FDis and species richness. As data from grid-cells is spatially non-independent, we fitted GLS models accounting for spatial autocorrelation in the error term using five correlation structures (exponential, Gaussian, linear, ratio, and spherical). This procedure resulted in 20 models for each group (Table S2). Then, we selected the best-fit model based on the Akaike information criterion (AIC) and on the spatial correlogram of the residuals (Figure S8). This analysis was performed in the *nlme* package. Subsequently, we included the four best-fitting GLS for each group in a SEM using the *piecewiseSEM* R package.

To analyse whether vertebrates have accumulated disparity in functional traits differently over time, we used disparity-through-time plots in the *geiger* R package. First, to describe the functional volume of each clade, we performed a Principal Coordinates Analysis onto the trait distance matrix (Figure S5). We chose only axes whose eigenvalues were higher than the Broken Stick criterion. We used a 95% confidence interval and ran 1,000 simulations to generate a null distribution of the disparity metric, calculated as average squared Euclidean distance.

## Results

### Spatial distribution

We found a congruent pattern of richness distribution among all groups in the central region of the Atlantic Forest, but not between frogs and Squamates (Table 1, Figure 2). We did not find a spatial congruence in FDis between ectotherms and between endotherms. However, we observed a high negative correlation between Squamates and birds. Specifically, squamates had higher FDis in the northern portion of the Atlantic Forest, while birds had higher FDis in the southern portion. We found higher FDis in the Serra do Mar for amphibians and in the innermost and central portions of the biome for mammals (Figure 1). We found a congruent pattern in diversification rate between mammals and ectotherms in the central region (Table 1, Figure 2). In contrast, the spatial distribution of MPD was not congruent among groups (Table 2). Ectotherm communities were older in the northern and younger in the southern portion. Bird communities had high MPD evenly distributed across the biome, with lower values in the central coastal region, and a cluster of greater FDis close to the Bahia endemism area and in the southern region. Mammalian communities were older in the central portion, specifically towards the interior of the continent (Figure 2).

**Figure 2.**
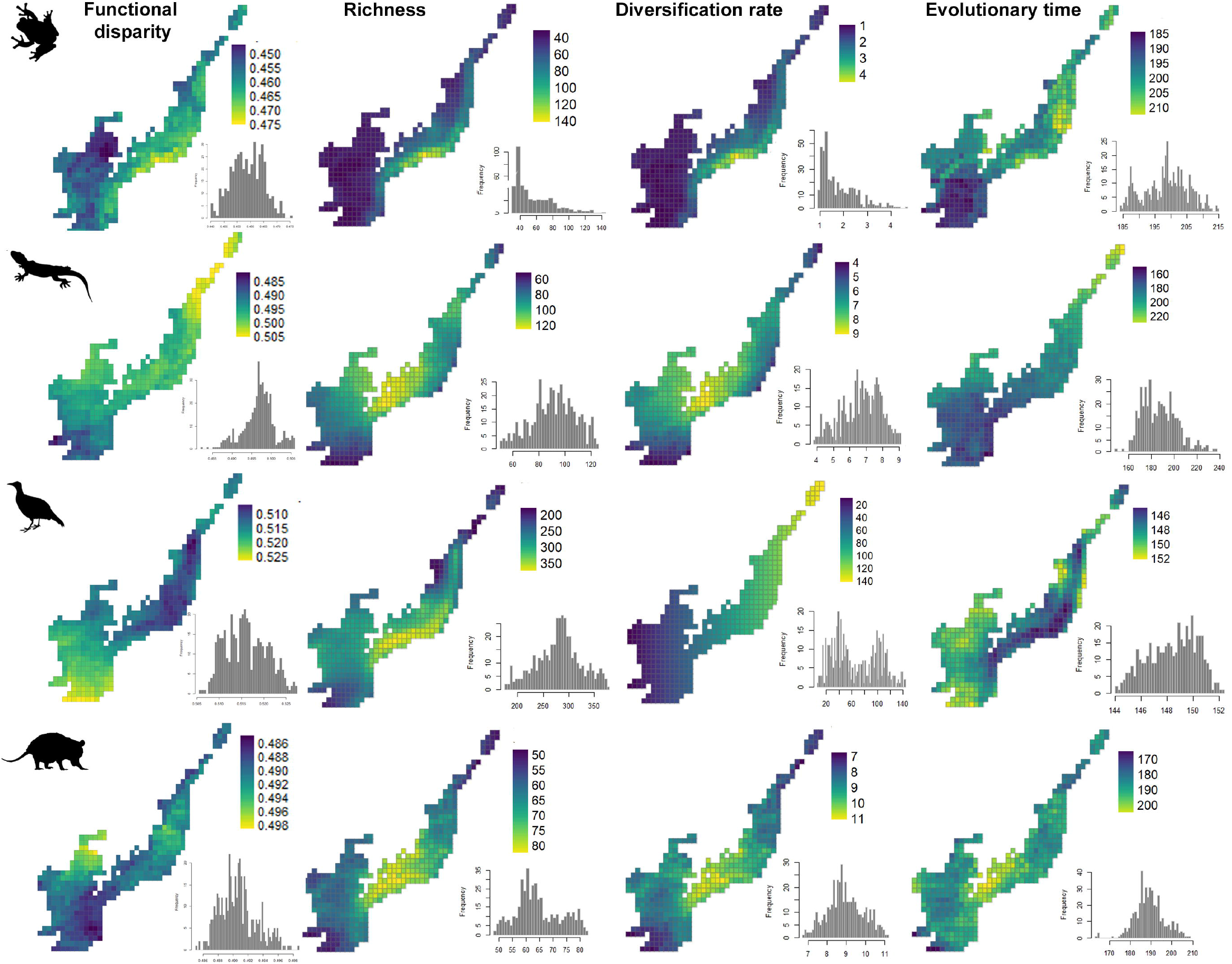
Geographical patterns in Functional dispersion (FDis), diversification rate (DR), and evolutionary time (MPD) of assemblages of frogs, Squamates, birds, and non-flying mammals of the Atlantic Forest. Silhouettes are CC-BY from PhyloPic.

**Table 1.**
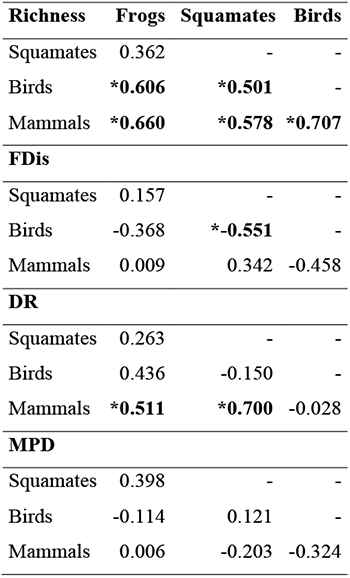
Pairwise spatial congruence of Richness, Functional Dispersion (FDis), Diversification Rate (DR) and Evolutionary time (MPD) between each clade of terrestrial vertebrates in the Atlantic Forest. Tjøstheim’s correlation coefficients above 0.46 are significant (*).

| **Richness** | **Frogs** | **Squamates** | **Birds** |
| --- | --- | --- | --- |
| Squamates | 0.362 | - | - |
| Birds | **\*0.606** | **\*0.501** | - |
| Mammals | **\*0.660** | **\*0.578** | **\*0.707** |
| **FDis** |  |  |  |
| Squamates | 0.157 | - | - |
| Birds | -0.368 | **\*-0.551** | - |
| Mammals | 0.009 | 0.342 | -0.458 |
| **DR** |  |  |  |
| Squamates | 0.263 | - | - |
| Birds | 0.436 | -0.150 | - |
| Mammals | **\*0.511** | **\*0.700** | -0.028 |
| **MPD** |  |  |  |
| Squamates | 0.398 | - | - |
| Birds | -0.114 | 0.121 | - |
| Mammals | 0.006 | -0.203 | -0.324 |

### Structural equation models

Non-equilibrium predictors were more important in explaining FDis and species richness of ectotherms and endotherms (Figure 3). Together, predictor variables better explained the variation in species richness of frogs, squamates, and mammals (R^2^= 0.99; R^2^=0.80; R^2^=0.70, respectively), than FDis (R^2^= 0.50; R^2^=0.50; R^2^=0.07, respectively). In contrast, predictors better explained the variation in FDis (R^2^=0.24), than species richness (R^2^=0.17) of birds.

**Figure 3.**
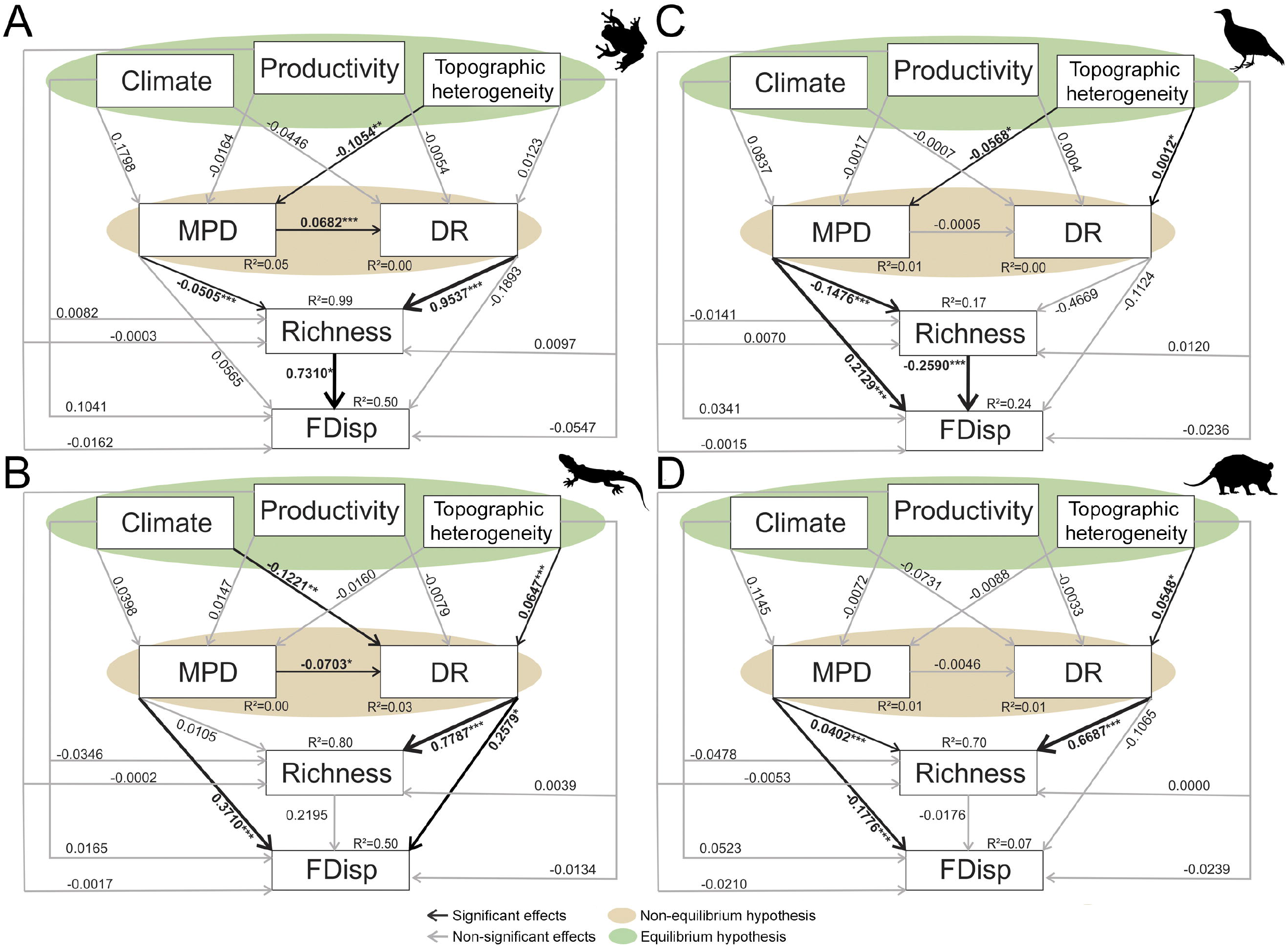
Path diagram showing the standardized coefficients (β) of models testing the influence of variables representing equilibrium hypothesis (Climate, productivity, elevation) and non-equilibrium (Mean Pairwise Distance [MPD] and Diversification Rate [DR]) on species richness and functional dispersion (FDis) of ectotherms: (A) Frogs and (B) squamates; and endotherms: (C) birds and (D) terrestrial mammals of the Atlantic Forest. Silhouettes are CC-BY from PhyloPic.

We found that MPD was a negative predictor of species richness in amphibians (coefficient = −0.050, *P* < 0.001) and birds (coefficient = −0.147, *P* < 0.001), and a positive predictor in mammals (coefficient= 0.402, *P* < 0.001). Diversification rates were positively related to species richness in mammals (coefficient= 0.6687, *P* < 0.001), amphibians (coefficient= 0.9537, *P* < 0.001), and squamates (coefficient= 0.7787, *P* < 0.001), but not birds (coefficient = −0.4669, *P* > 0.05). Assemblage age significantly influenced the FDis of squamates, birds and mammals as well as the species richness of frogs, birds, and mammals (Figure 3). Assemblage age was also correlated with diversification rate in ectotherms, but not endotherms. Diversification rates had the highest slopes for the species richness of frogs, squamates, and mammals. Conversely, at least one climate predictor was related to evolutionary variables and species richness in the four clades. Climate was related to diversification rates in squamates, and elevation was related to assemblage age in frogs (Figure 3). Elevation was related to diversification rate in squamates, birds and mammals.

## Discussion

Here, we used a functional biogeography approach to add new lines of evidence on ecological and evolutionary mechanisms involved in the diversification of species and traits in the Atlantic Forest. We found a significant and positive spatial pattern in the species richness among all groups, but only a weaker one between frogs and squamates. In contrast, the spatial distribution of trait diversity showed high negative values between birds and reptiles. Taken together, our results reinforce that time, trait evolution, and physiological constraints of ectotherms and endotherms are key drivers of vertebrate community assembly along the Atlantic Forest.

There is a complex geographic mosaic of evolutionary time, diversification rate, and trait diversity among vertebrates in the Atlantic Forest. One common pattern is that the central region of the biome has the highest species richness for all clades. Yet while richness was mostly congruent between groups, squamates and frogs had different patterns, with greater squamate diversity at the border with dry biomes (Figueiredo *et al.*, 2021), while frogs had greater richness in coastal areas with rainforest that experiences high annual precipitation and low seasonality (Benício *et al.*, 2021). This difference can be explained by their distinct physiological requirements and the water dependence of frogs (Lourenço-de-Morais et al. 2020). Despite clear spatial structuring of species diversity across the Atlantic Forest region, richness was most strongly predicted by diversification rates in frogs, mammals, and squamates as predicted by a non-equilibrium model, rather than directly by climate as predicted under an equilibrium model. The evolutionary speed hypothesis predicts that higher temperatures support faster speciation rates (Rohde 1992). Ectotherms, which regulate their metabolism based on environmental temperatures, should have a greater climate dependency on speciation rates than endotherms (Gillooly & Allen 2007). Further, elevation played an indirect role on DR in birds, mammals, and squamates, but not frogs, as recently found (García-Rodríguez et al. 2021), which might be explained by mountains acting as speciation pumps in those groups. Conversely, elevation was related to assemblage age in frogs and birds. The central Atlantic Forest encompasses the Serra do Mar range that experienced climatic stability during the Last Glacial Maximum. This mountainous area, along with past climatic variation, has contributed to increase diversification rates by playing a key role in allopatric and ecological speciation, decreasing extinction rates, and constraining species dispersal (Rahbek et al., 2019, Benício *et al.*, 2021).

Although richness of the four groups was concentrated in the central Atlantic Forest, FDis had idiosyncratic patterns among taxa. The areas of greatest FDis are in ecotones; the Atlantic Forest and Caatinga transition for squamates, regions close to the Pampas in birds, and in the transition to the central region of Cerrado in mammals. For frogs, we found high values of functional dispersion in the Serra do Mar, a region with highland grasslands, but known for harbouring a high species richness. This is in agreement with our findings, where richness is a strong predictor of frog functional dispersion. Likewise, recent studies suggest that the rate of diversification and the time for speciation explain why there are more frog species in the coastal region than in the interior region (Benicio *et al.*, 2021). Ecotones might support high FDis because they typically have high habitat heterogeneity, which permits a high degree of niche partitioning and co-occurrence of functionally distinct species (Smith et al. 2001). Ecotones are also harbour high degrees of endemism and uniquely adapted species (Kark et al. 2007). One reason why ecotone harbour high biodiversity is that they drive faster rates of ecological speciation along environmental gradients or among habitat patches due to higher heterogeneity (Schiltzhuizen 2000). However, we did not find any significant relationships between FDis and DR. Instead, the strong positive association between MPD and FDis suggests that functionally diverse ecotone regions have formed from (1) a greater evolutionary time available for functionally distinct lineages to accumulate, (2) more time for the disparification of traits to evolve *in-situ.* This is the opposite for mammals, in which MPD and FDis were negatively related, meaning that recently occupied grid-cells have more trait diversity. We used MPD as a proxy for evolutionary time (Oliveira et al 2016), but it could reflect the more recent assembly of phylogenetically distant taxa. Although difficult, further studies may focus on modelling the colonization dynamics that occurred between bioregions and separating these two scenarios.

Different drivers of species richness, and FDis mean these two variables are decoupled in the Atlantic Forests. Functional dispersion shows no relationship with species richness in squamates, and mammals, while in birds and frogs FDis was significantly, but negatively related to species richness in birds, and positively in frogs. This pattern indicates that while species-rich communities of birds had functionally redundant species, while in frogs it is the opposite, with greater richness and greater trait diversity. Therefore, as bird species entered communities, they more likely were closer to the centroid in trait-space (Kuebbing *et al.*, 2018). Decoupled FDis and richness supports the hypothesis that separate processes generate these two variables across vertebrates (Oliveira *et al.*, 2016). Furthermore, this result suggests that, not only do these factors not positively covary, but also that high diversity can exist with high functional redundancy, which has also been shown in several global studies of lizards (Skeels et al. 2019), birds, and mammals (Cooke et al 2019). One prediction is that increases in species richness should lead to positive, but non-linear and asymptotic increases in trait diversity (Halpern & Floeter, 2008). However, while functional richness and species richness are expected to covary, FDis, which is a measure of how divergent species are from each other in trait space, may not always follow the same pattern (e.g., MacDougall *et al.*, 2019).

The northern Atlantic Forest had the highest FDis for squamates and oldest communities for ectotherms, but they were not congruent. In contrast, birds had an opposite pattern, with greater FDis and older communities in the south. Climatic oscillations and the Pleistocene arid phases were greater in the southern Atlantic Forest (Carnaval, Hickerson, Haddad, Rodrigues, & Moritz, 2009). Therefore, this pattern may be related to the different physiological constraints, dispersal ability, and colonization time between ectotherms and endotherms towards this region (Araújo & Pearson, 2005). The northern region had past connections with the Amazon that promoted intraspecific diversity of both frogs and squamates (reviewed in Peres et al., 2020). Due to the high speciation rates in the Amazon rainforest, several taxa later dispersed to other regions (Antonelli *et al.*, 2018). For example, amphibians have narrow physiological requirements (thermoregulation and evapotranspiration) making areas with lower latitudes and high altitudes harbour smaller-sized species (Vasconcelos *et al.*, 2019; Lourenço-de-Moraes *et al.*, 2020). Conversely, endotherms are more tolerant of climate change, which could have allowed birds to colonize the southern region earlier or more often than ectotherms. Due to their greater dispersal ability and broader thermal tolerance, birds may have used different colonization routes than ectotherms (Andes-South of Atlantic Forest; Nores, 2020). Thus, past glacial cycles that connected tropical forests in South America and allowed the exchange of species, plus the colonization and dispersal ability of each group, may have been the main drivers of the trait diversity of Atlantic Forest vertebrates.

Non-equilibrium dynamics, including greater evolutionary time and faster diversification rates, were relatively more important in explaining the species richness and FDis in all groups. Older communities accumulated higher FDis, especially in ectotherms, while species-rich communities had higher diversification rates. Previous studies found that non-equilibrium dynamics are important in explaining species richness at global scales. For example, evolutionary time has repeatedly been the strongest predictor of richness in tetrapods (Marin *et al.*, 2018) and fish (Miller & Román-Palacios, 2021). Conversely, evidence that species richness is driven by faster diversification rates is mixed, with many recent studies reporting faster diversification rates in regions with fewer species (Weir & Schluter 2007; Rabosky et al. 2015; 2018). In contrast, DR, but not evolutionary time, was a strong predictor of species richness. However, evolutionary time a better predictor of FDis. A potential explanation for this discrepancy is the spatial scale of analysis, as our study excluded some high-latitude areas found in those studies to have the highest rates. Regardless, we find little evidence for a direct role of equilibrium dynamics. Climate, productivity, and elevation are all expected to increase resource availability or the diversity of resources that support more species. In some cases, these variables were strongly related to response variables via assemblage age and DR. Equilibrium dynamics could influence DR by increasing competition between species, driving ecological speciation (Skeels & Cardillo 2019), or by increasing population sizes and genetic diversity (Gillman & Wright 2013). As such, the “ecological limits” hypothesis cannot be excluded as an indirect explanation for richness patterns (e.g., Mittelbach *et al.*, 2007; Harmon & Harrison, 2015), but is an unlikely explanation for patterns of FDis, which showed no direct relationships with climate or elevation, and only a single indirect relationship via MPD in birds. Unfortunately, trait data are not evenly distributed across clades, and may be lacking for some species with smaller body size, for example (Jones *et al.*, 2009; Johnson *et al.*, 2020). Further studies should include more species with trait and phylogeny data to confirm our findings.

Our results are in line with the idea that continental communities are temporary set of species in ever-changing environments (Jackson & Overpeck, 2000; Harmon & Harrison, 2015). Species respond to abiotic and biotic filters, which generate differences in diversification rates between clades over time (Harmon & Harrison, 2015). Ecological opportunities in newly formed environments, such as those in the Atlantic Forest in the past, can produce high speciation rates (Antonelli *et al.*, 2018). An adaptive radiation can occur following the colonization of new habitats, extinction of competitors or the raise of key evolutionary innovations that allow the invasion of new adaptive zones (Heard & Hauser, 1995; Yoder *et al.*, 2010). Intraspecific data (Peres et al. 2020) and climate modelling (Carnaval et al. 2009) show that the biotas of the Atlantic Forest were strongly influenced by Plio-Pleistocene glacial cycles, altering the ranges of organisms, and promoting biogeographic links between forested domains during different climatic phases (Sobral-Souza *et al.*, 2015). Areas of greater climate stability may harbour older communities that accumulate greater trait diversity over time (Oliveira *et al.*, 2016), as observed here. Therefore, community assembly over time more likely takes place without defined ecological limits at continental scales (Harmon & Harrison, 2015; Oliveira *et al.*, 2016), with climatic variables playing a secondary role.

Here, we added new evidence on ecological and evolutionary mechanisms to the diversification of vertebrates and their traits in the Atlantic Forest. Non-equilibrium mechanisms played a primary role in driving species richness and FDis, with elevation and climate variables having an indirect effect. Diversification rates were more related to richness, while assemblage age was more related to FDis, except for mammals, in which FDis and MPD were negatively related. It is possible that areas of greater climate stability in the past may have served as refuge and harbour older communities, which subsequently accumulated greater trait disparity and richness over time (Oliveira et al. 2016). Furthermore, we found a congruent pattern of species richness among all groups, except among ectotherms. In contrast, for other metrics such as evolutionary time and trait diversity, we found idiosyncratic and distinct patterns for each group. Our results suggest that the time of colonization and the exchange of species between past tropical forests may have acted directly in the assembly of Atlantic Forest communities.

## Acknowledgment

Gabriel Tirintan helped collecting bird data and and Mario R. Moura provided data for Squamata.

## Author’s contribution statement

DBP and FRS conceived the study. MTM obtained all data, conducted data analysis, and wrote the first version of the manuscript. AS contributed to the final version of the text and gave input on figure presentation and data analysis.

**Biosketch** — Matheus de T. Moroti is interested in natural history in the broadest sense, investigating population dynamics, community ecology and biogeography.

## Data Availability Statement

All data and associated R scripts are available at https://doi.org/10.6084/m9.figshare.19735552.v1

